# A comparative analysis of the antiviral response in two bat species reveals conserved and divergent innate immune pathways

**DOI:** 10.1101/2023.04.23.537989

**Authors:** Lilach Schneor, Stefan Kaltenbach, Sivan Fridman, Yomiran Nissan, Gal Shuler, Evgeny Fraimovitch, Aleksandra A. Kolodziejczyk, Maya Weinberg, Giacomo Donati, Emma C. Teeling, Yossi Yovel, Tzachi Hagai

**Affiliations:** Shmunis School of Biomedicine and Cancer Research, George S Wise Faculty of Life Sciences, Tel Aviv University, Tel Aviv 69978, Israel; School of Zoology, George S. Wise Faculty of Life Sciences, Tel Aviv University, 6997801, Tel Aviv, Israel; International Institute of Molecular and Cellular Biology, Warsaw, Poland; Department of Life sciences and Systems Biology, University of Turin, Torino, Italy; Molecular Biotechnology Center, University of Turin, Torino, Italy; School of Biology and Environmental Science, University College Dublin, Dublin, Ireland; Sagol School of Neuroscience, Tel Aviv University, 6997801, Tel Aviv, Israel

## Abstract

Bats host a range of viruses that cause severe disease in humans without displaying clinical symptoms to these infections. The mechanisms of bat adaptation to these viruses are a continuous source of interest but remain largely unknown. To understand the landscape of bat antiviral response in a comprehensive and comparative manner, we studied this response in two bat species - the Egyptian fruit bat and the insectivore Kuhl’s pipistrelle, representing the two major bat subordinal clades. We profiled the transcriptional response to dsRNA – that triggers a rapid innate immune response – in skin fibroblasts from a large cohort of replicates from each bat species, using RNA-sequencing, and compared bat response with responses in primates and rodents. Both bat species upregulate a similar set of genes, many of which are known to be involved in the antiviral response across mammals. However, a subset of these genes is transcriptionally divergent in response between the two bat species.

These transcriptionally divergent genes also evolve rapidly in coding sequence across the bat clade and have particular regulatory and functional characteristics, including specific promoter architectures and association with expression programs thought to underlie tolerance and resistance in response to viral infection. In addition, using single-cell transcriptomics, we show that transcriptionally divergent genes display high expression variability between individual cells. A focused analysis of dsRNA-sensing pathways further points to significant differences between bat and human in basal expression of genes important for triggering antiviral responses. Finally, a survey of genes recently lost or duplicated in bats points to a limited set of antiviral genes that have undergone rapid gene loss or gain in bats, with the latter group resulting in paralogs displaying divergence in both coding sequence and expression in bat tissues. Our study reveals a largely conserved regulatory program of genes upregulated in response to viral infection across bats and other mammals, and points to a set of genes that evolved rapidly in bats through multiple evolutionary mechanisms. This divergence can contribute to bat adaptation to viral infection and provides directions to understanding the mechanisms behind it.

## Introduction

Among mammals, bats display unique life histories and adaptations, including powered flight, extreme longevity relatively to their body mass and echolocation. In addition, bats have been shown to display subclinical symptoms when infected with several viruses that can cause severe disease in humans^1–4^. In some of these cases, bats were confirmed to be the natural reservoirs of these viruses^1–5^. For example, the Egyptian fruit bat, *Rousettus aegyptiacus*, is thought to serve as a reservoir of the highly pathogenic Marburg virus^6^, based on field collection from wild bats^7–9^ and on a series of ecological and experimental studies^8, 10–15^. In addition, infection of *Rousettus* with the related Ebola virus is asymptomatic, although the exact reservoir species is not yet known^12, 16, 17^.

Several coronaviruses – severe acute respiratory syndrome coronavirus (SARS-CoV), SARS-CoV-2 and Middle East respiratory coronavirus (MERS-CoV) – that recently transferred to humans, are also thought to originate in bats, although other mammals likely acted as proximate reservoirs for human infection^18–21^. Furthermore, direct and indirect bat-to-human spillovers of two henipaviruses – Nipah virus and Hendra virus – lead to infections with high mortality rates in humans^22–24^, leading to concerns of potential future pandemics from henipaviruses. The unique lifestyles of bats, including flight, rapid changes in body temperature and crowded colonies in closed environments such as caves, were all suggested to facilitate bat adaptation to these viruses^2^.

The links between bat and human pathogens and the often asymptomatic or mild infections observed in bats, have led to a great interest in bat viriome and immunity, and to the notion that bats disproportionately contribute to emerging zoonotic viruses that have crossed to humans^25^. However, a recent survey of virus-reservoir relationships challenged this notion, when controlling for the size of the bat clade^26^. Bats are the second largest mammalian order after rodents, comprising ∼22% of known mammalian species, and have adapted to diverse ecological niches across the planet^27, 28^. Thus, the clade’s size and large diversity as well as bat occurrence across the globe, have contributed to facilitating zoonotic transfers of bat viruses to humans and other mammals.

Regardless of this, a series of studies focusing on specific bat species, immune pathways and viral infections revealed unique adaptions in the bat immune system. These include the constitutive expression in uninfected cells and tissues of the black flying fox (*Pteropus alecto*) of IFNα – an antiviral cytokine that is usually only upregulated following infection^29^. Other innate immune-related genes, including known interferon-stimulated genes, were shown to be highly expressed in uninfected cells of several bat species^30–32^. High expression of various innate immune genes may point to greater resistance to viral disease, by rapid inhibition of infecting viruses^5, 33^. In contrast, tolerance to disease – a mechanism by which viruses and other pathogens replicate in host cells without leading to an excessive immune response and thereby decreasing tissue damage, was also suggested to be related to mild symptomatic infection of certain bat species^1, 5, 15^. For instance, bats were shown to have a dampened inflammasome activation in comparison with other mammals through various mechanisms, including the loss of the PYHIN gene family locus that encodes for inflammasome DNA sensors^34^, and the reduction of inflammation and apoptosis mediated by the inflammasome sensor NLRP3^35^. In addition to these mechanisms, certain antiviral and immune gene families were shown to expand or to contract in bats or in specific branches of the bat clade and to display signatures of positive selection in their coding sequences when compared between bat species, including type-I interferons and important antiviral proteins APOBEC3^29, 36, 37^, PKR^38^ and Tetherin^39^.

Here, we chart the transcriptional landscape of the antiviral innate immune response in a comparative framework, by triggering this response in primary cells from two bat species. We focus on representatives of the major bat subordinal clades^40^ - the Egyptian fruit bat, *Rousettus aegyptiacus,* and the insectivore bat *Pipistrellus kuhlii* (Kuhl’s pipistrelle). Various aspects of *Rousettus* innate and adaptive immunity have been studied^13, 41–46^, due to its asymptomatic infection of filoviruses, as well as its association with infection with other viruses^47, 48^. In contrast, antiviral immunity of *Pipistrellus*, a distantly related bat to *Rousettus*, remains poorly characterized, despite studies that suggested the presence of several Alpha- and Beta-coronaviruses (including closely related variants to MERS-CoV) in its population in Europe and the Middle East^49–54^. The fact that *Pipistrellus* is common in agricultural and urban areas and can be in close contact with humans, underlines the importance of studying its viriome and immune system.

We triggered the innate immune response – an expression program that involves the upregulation of cytokines and chemokines, restriction factors that inhibit viral replication and gene related to apoptosis and to the regulation of this response^57, 58^, in skin fibroblasts from both bat species. Fibroblasts play key roles in infected tissues^59, 60^, and, due to their robust response, are often used as experimental models to study the innate immune response across species^56, 61, 62^ and between human individuals^63, 64^. We profiled the transcriptional response in both species and employed a comparative genomics approach to find transcriptionally conserved and divergent innate immune genes between the two bat species. We then analyzed the regulatory and functional characteristics of these conserved and divergent genes, and compared the results in bats with analogous results in primates and rodents. We also studied innate immune gene upregulation in individual bat and human cells using single-cell RNA-seq. Finally, we studied the basal level of expression of these innate immune genes in various tissues in human, mouse and bat as well as the transcriptional and coding sequence divergence between innate immune genes that have duplicated in bats.

Our study thus maps the evolutionary landscape of bat innate immune response, links functional and regulatory features of innate immune genes with their transcriptional and coding sequence evolution, and points to innate immune genes that diverged in bats – divergence that may be play a role in their adaptation to viral infection.

## Results

### An *in vitro* stimulation system to characterize bat antiviral response

To study the evolution of the antiviral innate immune response program we grew skin fibroblasts from wing biopsies of two bat species - the Egyptian fruit bat, *Rousettus aegyptiacus,* and the insectivore bat *Pipistrellus kuhlii* (Kuhl’s pipistrelle). We triggered an antiviral response using polyinosinic:polycytidylic acid (poly(I:C)), a synthetic double-stranded RNA (dsRNA) sensed by various sensors, such as TLR3, MDA5 and IFIH1. Unlike viral infection that can lead to a virus-specific response and where the virus can antagonize the host immune system, dsRNA leads to a general antiviral response, unmodulated by viruses. We profiled the resulting transcriptional changes using RNA-seq from 10-12 biological replicates (*Rousettus*: 6 females and 6 males; *Pipistrellus*: 3 females and 2 males, each in two biological replicates) between stimulation and control (see **Figure 1A** and **Supporting Table 1** for a detailed list of samples). We used the same stimulation conditions (in terms of time and concentration of dsRNA transfection) as we did previously, with an analogous fibroblast system from primates and rodents^56^. For analysis of cell-to-cell variability in gene expression before and after stimulation, we also profiled transcriptomes of single cells from *Rousettus* and human, as detailed in the relevant sections below.

**Figure 1:**
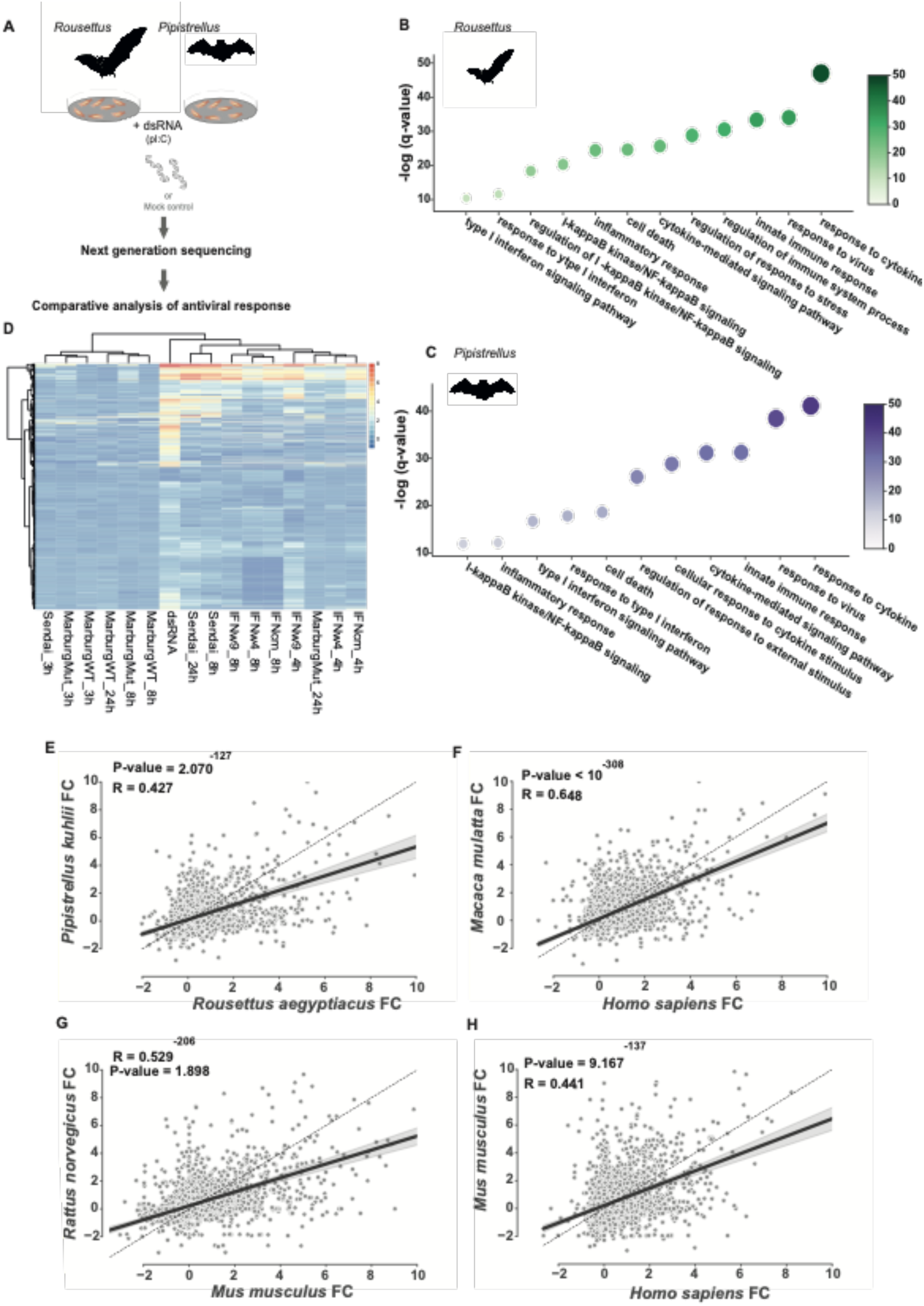
Characterization of dsRNA-stimulated genes in of cells from *Rousettus* and *Pipistrellus*. (A) System overview: dermal fibroblasts derived from wing biopsies from two bat species were cultured and stimulated with dsRNA and control, followed by profiling of the response using bulk RNA-seq, gene quantification and differential expression analysis using edgeR. **(B-C)** Go term enrichment analysis of genes upregulated in response to dsRNA-stimulus in (B) *Rousettus* and (C) *Pipistrellus*. Selected non-redundant terms are shown in a decreasing order of significance (FDR-corrected P-values are shown). Detailed analyses, including all enriched terms appear in Supporting Tables S4 and S5, respectively. **(D)** Heatmap of log2(fold change) of dsRNA-stimulated *Rousettus* genes across different immune stimuli with different types of IFN and in response to infection with Sendai virus and Marburg virus. **(E-H)** Spearman’s rank correlation of gene fold change in response to dsRNA, in 2,865 one-to-one orthologous genes between (E) *Rousettus* and *Pipistrellus*, (F) Human and Macaque, (G) Mouse and Rat, and (H) Human and Mouse. Genes included are dsRNA-regulated (FDR-corrected<0.01) in at least one of the six species appearing in figures E-H.

### DsRNA-regulated bat genes are enriched with innate immune functions

We first studied the transcriptional response of fibroblasts to stimulation in each of the bat species separately by performing differential expression analysis between control and stimulation using edgeR^65^. We observed that 968 and 840 genes in *Rousettus* and *Pipistrellus*, respectively, are upregulated in response to stimulus (logFC>0 and FDR-corrected P-value<0.01)), while significantly fewer genes are downregulated (logFC<0 and FDR-corrected P-value<0.01) - 61 and 81 genes in the respective species. Both numbers and fractions of differentially up- and down-regulated genes from the overall expressed genes are similar to those observed in analogous dsRNA stimulations of dermal fibroblast cells in primates and rodents^56^. For example, in dsRNA-stimulation of human fibroblasts, there are 1,255 and 161 upregulated and downregulated genes, respectively (see detailed list of bat genes with their DE values in **Supporting Tables 2-3**). In both bat species we observe a strong upregulation of antiviral and inflammatory cytokines and chemokines, including IFNB, CXCL10 and CCL5, as well as other genes involved in the antiviral response, such as transcriptional factors and signal transducers - IRFs and STATs, and various restriction factors, including SAMHD1 and ADAR.

We next used g:Profiler^66^ to characterize pathways enriched in the set of upregulated genes in each bat species. We observed that in both bats the upregulated genes are involved in pathways known to be related to the primary and secondary waves of the antiviral response^56^, including terms such as ‘response to virus’ and ‘response to cytokine’, as well as other associated processes such as ‘cell death’ and ‘inflammatory response’ (see a set of non-redundant terms for each species in **Figure 1B-C** and the full list of enriched terms in **Supporting Tables 4-5**). These pathways are similar to those observed to be enriched in human and mouse dsRNA stimulation of fibroblasts^56, 62, 64^.

To further compare the genes upregulated in our dsRNA-stimulation system with other related systems, we used available data of a Rousettus cell line (RoNi/7.1 - immortalized kidney cells), where cells were either stimulated with interferons (IFNs) or infected with wildtype or mutant strains of RNA viruses (see Figure 1D, Table S6 for full list of datasets and Table S7 for gene expression values)10,42. The transcriptional profile of response to dsRNA from our experiments clusters with Sendai virus infection and with stimulations with the set of IFNs. Late Sendai virus infections (8 and 24 hours) were more similar to dsRNA-stimulation (of 4 hours) in comparison with earlier infection (2 hours), likely reflecting differences in kinetics of RNA recognition between the two systems. Interestingly, wildtype Marburg virus infections cluster separately from dsRNA- and IFN-stimulation as well as from most Sendai virus infections.

We next looked at gene expression across individual cells within the fibroblast population using the single-cell transcriptomics data of human and *Rousettus*. For this, we profiled single-cell transcriptome of thousands of *Rousettus* and human cells, with and without dsRNA stimulation using Chromium Single Cell 3′ gene expression v.3.1 (see **Methods**). We observe that the regulatory architecture of cytokine expression previously reported by us and others^67–71^ is recapitulated in our human and *Rousettus* cells: several types of cytokines, such as IFNs, are only expressed in a few stimulated cells in both species, whereas others, such as chemokines from the CXCL family, are more widely expressed (**Supporting Figure 1**).

Finally, we tested whether the use of skin samples from different regions in the body may affect our results. We observed that the upregulated genes and their level of upregulation are highly similar between fibroblasts derived from two different skin regions in *Rousettus* (P-value<10^-293^, **Supporting Figure 2**).

Together, our above analyses suggest that dsRNA-stimulation of bat dermal fibroblasts leads to a strong upregulation of genes involved in conserved antiviral pathways, similar to other mammalian species, including primates and rodents.

### Transcriptional response divergence between bat species and between bats and other mammals

We next focused on the overall similarity in transcriptional response to dsRNA between species, by comparing the response level to dsRNA between orthologous genes across bats and other mammals. For this, we first inferred the set of orthologs between the two bat species as well as between these bats, two primate species (human and macaque) and two rodent species (mouse and rat), using the eggNOG program^72^ (see analysis details in **Methods**). The resulting gene orthology table between the two bats is available as **Supporting Table S8,** and the one-to-one orthology table for the six species is available as **Supporting Table S9**.

We focused on a set of 2,865 one-to-one orthologous genes that are differentially expressed in response to dsRNA in at least one of six mammalian species (the two bat species from our current analysis, human, rhesus macaque, mouse and rat). We performed a correlation analysis between their fold change in response to dsRNA stimulation (see differential expression results based on edgeR^65^ analysis for all 6 species in **Table S9**). We observed a significant correlation between the two bat species (**Figure 1E**, Spearman’s rank correlation ρ=0.427, P-value<10^−27^), corresponding to an overall similar response between the two bat species. However, when performing the correlation analysis on the same set of genes using a similar fibroblast dsRNA-stimulation system in primates and rodents (human versus macaque; mouse versus rat) we observed a higher correlation in transcriptional response between the two primates as well as between the two rodents (**Figures 1F-G**, ρ=0.648 and 0.529, respectively). When comparing human versus mouse transcriptional response, we observe a similar correlation to that observed between the two bat species (ρ=0.441 between human and mouse versus ρ=0.427 between the two bats). This seemingly low correlation between the two bat species is in agreement with the fact that these two bat species belong to the two major bat suborders, Yangochiroptera and Yinpterochiroptera, whose last common ancestor is predicted to have existed ∼60 million years ago^40, 73^ (while primates and rodents split approximately 90 million years ago^74, 75^). This finding is consistent with the notion that different bat species, while having a largely conserved genetic program upregulated in response to viral infection, may still differ in the level of upregulation of specific genes and that there may be high diversity across bat species in their response to viruses. The characteristics of the most divergent antiviral genes between the two studied bat species are the focus of the subsequent sections of our analysis.

### Transcriptionally divergent bat innate immune genes display specific evolutionary and regulatory characteristics

To study the characteristics of transcriptionally conserved versus divergent bat antiviral genes, we focused on the same set of 2,865 differentially expressed genes from the previous analysis. These genes were partitioned into three equally sized groups, displaying high, medium and low levels of transcriptional response divergence in response to dsRNA between the two bat species (see **Methods** for details on the metrics used to calculate gene’s divergence level. The divergence scores are found in **Table S7**).

We first compared the rate at which the genes in the three groups evolved in coding sequence. Thus, we asked whether antiviral genes with different levels of transcriptional divergence in response to dsRNA, also evolved at different rates also at the coding sequence evolution. This was done (1) by considering the ratio of substitution rates at non-synonymous and synonymous sites (dN/dS) in orthologs from a set of 18 bat species^76^ (**Figure 2A**), and (2) by comparing the sequence similarity between the orthologs of the two bat species (**Figure 2B**) (per-gene values are available in **Table S7**). In both measures, we observe that innate immune genes with transcriptionally divergent response tend to have higher coding sequence divergence than the group of innate immune genes with conserved response (**Figure 2A-B**, P-value = 0.017 and 1.5×10-16, respectively, one sided Mann-Whitney test).

**Figure 2:**
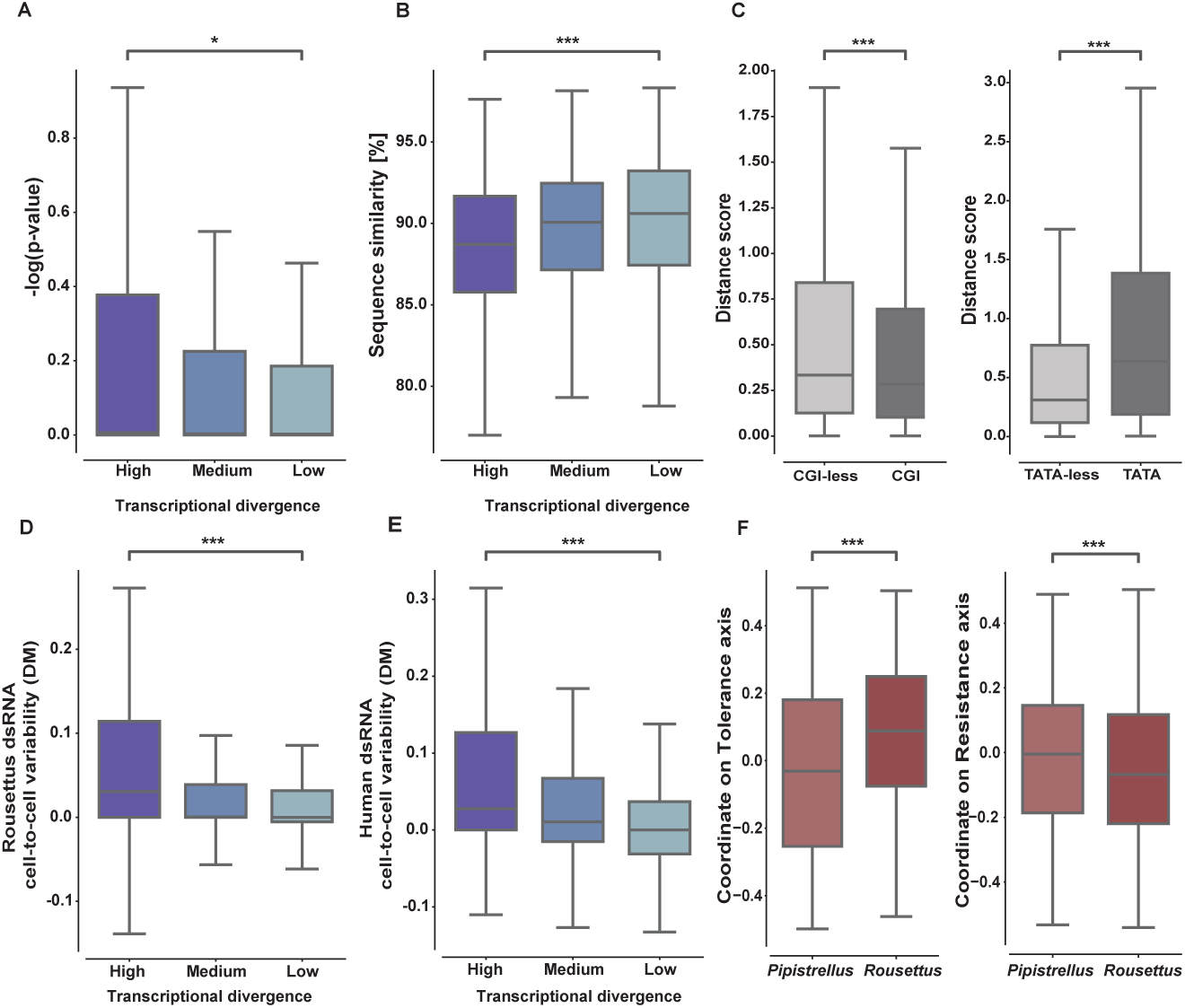
Regulatory and evolutionary characteristics of conserved and divergent dsRNA-stimulated genes between *Rousettus* and *Pipistrellus*. (A-B) Coding sequence divergence *versus* transcriptional divergence: Genes were partitioned into three groups based on divergence in response to dsRNA stimulation between the two bat species (termed high, medium and low transcriptional divergence), and coding sequence divergence was compared between these three groups: (A) Distributions of P-values of test for positive selection across bat orthologs (using the dN/dS values from a previous analysis^79^), and (B) sequence similarity (precent of identity) between the two bat species, are shown for the three gene groups (1,899 and 2,728 genes with available data in A and B, respectively). High- and low-divergence groups are compared using a Mann-Whitney test. (C) Divergence in response to dsRNA between the bat species in genes with and without CpG Island (CGI) in their promoters, and with and without TATA-box in their promoters (896 CGI and 1,969 CGI-less genes; 107 TATA and 2,758 TATA-less genes). Groups are compared using a Mann-Whitney test. (D-E) Using single-cell RNA-seq, the distribution of cell-to-cell variability in gene expression, as measured using the distance to median approach (DM), is shown for each of the three groups mentioned in (A), for (D) dsRNA-stimulated *Rousettus* cells and (E) stimulated human cells (1,980 and 2,578 genes with computed DM values, respectively). High- and low-divergence groups are compared using a Mann-Whitney test. See Supp Fig 3 for analysis of unstimulated *Rousettus* and human cells. (F) The distribution of measures for tolerance and resistance are shown for genes that are highly upregulated in response to dsRNA in *Rousettus* versus *Pipistrellus*, or the opposite (1,928 and 1,949 genes, respectively). The distribution of *Rousettus*-specific and *Pipistrellus*-specific genes are comparted using a Mann-Whitney test. (* = P<0.05, *** = P<0.001).

Previous work suggested that gene variability in expression between species (as well as between different conditions) is associated with different promoter architectures^56, 77^: CpG Islands (CGIs), a major element in core promoters of around half the mammalian genes, are thought to be associated with homogenous transcription between conditions and between species of nearby genes. In contrast, TATA-box elements that are relatively rare in mammalian promoters, are associated with variable transcription of their regulated genes^78^. We assessed the presence of both elements (available in **Table S7**) in core promoter regions of *Rousettus* genes (see **Methods**), and observed that genes with CGI promoters are significantly more conserved in their transcriptional response to stimulus than genes that lack CGIs in their promoters. In contrast, genes with TATA-box elements in their promoters exhibit greater transcriptional response divergence than those genes without TATA-box (**Figure 2C**, P-value<0.001 in both comparisons, one sided Mann-Whitney test).

We next studied whether gene transcriptional response divergence between species is linked with gene expression variability across individual cells from the same species. For each gene expressed in the single-cell data that is upregulated in response to dsRNA (based on our differential expression analysis above), we estimated cell-to-cell variability in expression across individual cells in a manner that takes into account their mean expression level, using the distance to median (DM) approach^79^. This was done separately for stimulated and unstimulated cells, and for both human and *Rousettus* cells (data available in **Table S7**). We observed that in all tested sets of cells, the group of genes displaying high transcriptional divergence between the two bat species also displays high cell-to-cell variability in expression (high DM values), which is significantly higher than the variability observed in the group of low divergence between species (low DM values) (P-value<0.001 in both cases, one sided Mann-Whitney test, **Figures 2D-E** and **Supporting Figure 3**).

Thus, innate immune genes diverging in transcriptional response between bat species tend to vary between individual cells within the same species, and have also undergone rapid evolution at the coding sequence level compared with innate immune that display conservation in their transcriptional response between bat species.

### Characteristics of *Rousettus*-specific and *Pipistrellus*-specific antiviral genes

So far, we compared bat antiviral genes with high, medium and low levels of transcriptional divergence in response to dsRNA between the two bat species, using a continuous divergence measure. Next, we asked whether there are defining characteristics for genes significantly more upregulated in one of the two species with respect to the other bat species - i.e., *Rousettus*- and *Pipistrellus*-specifically upregulated genes. For this, we used the top 20% of genes (2,433 genes out of all differentially expressed genes appearing as one-to-one orthologs in both bat species) that are most highly upregulated in *Rousettus* with respect to *Pipistrellus*, as well as the opposite group, essentially taking the two extremes of the group of high-divergence level. These two subgroups will be termed for simplicity *Rousettus-* and *Pipistrellus*-specific genes (see **Methods** for the detailed procedure).

By looking at enriched pathways using g:Profiler^66^ and by contrasting *Rousettus-* and *Pipistrellus*-specific genes, we observe that *Rousettus*-specific genes are associated with various developmental processes, including ‘anatomical structure morphogenesis’ and ‘tissue development’. *Pipistrellus*-specific genes are associated with ‘Wnt signalling pathway’ and ‘lipid metabolic process’ and various processes associated with protein degradation including ‘regulation of proteolysis’ and ‘deubqiuitination’ (see **Tables S10-S15**). The enrichment of the above-mentioned pathways that are not “core antiviral pathways” is consistent with the notion that significant evolutionary changes would occur in a large number of genes belonging to pathway at the “periphery” of the antiviral response.

We next asked whether these *Rousettus-* and *Pipistrellus*-specific genes are associated with expression programs that underlie “disease tolerance” (containment of infection while avoiding an excessive immune reaction) and “disease resistance” (effective inhibition of infection). Previous works suggested that different species of bats may display greater disease tolerance or greater resistance to virus infections with respect to other mammals due to their unique lifestyle^5, 15^. The extent of tolerance and resistance likely differs between various bat species as well as depending on the specific pathogen in question. A recent study used an array of mouse strains infected with influenza virus and profiled both the physiological response of the mice and their gene expression during infection. This has yielded distinct sets of genes that are associated with greater tolerance or resistance to infection^80^. These sets were shown to be indicative of tolerance or resistance in a spectrum of infectious diseases and conserved across species. We compared the set of *Rousettus-* and *Pipistrellus*-specific genes in terms of their correlation with resistance- and tolerance-associated programs (**Figure 2F**, additional results are found in **Supporting Figure S4**, detailed data is in **Table S16**). Interestingly, we observed that the transcriptional program generally associated with disease tolerance is significantly more prominent in *Rousettus*, while the molecular program primarily associated with resistance is significantly more active in *Pipistrellus* (P-value<0.001, one sided Mann-Whitney test, in both comparisons, see **Figure 2F**).

In summary, genes displaying divergence in their level of upregulation between the *Rousettus* and *Pipistrellus*, are enriched with different cellular pathways that do not represent core antiviral pathways. Furthermore, the two species differ in the activities of expression programs associated with disease tolerance or resistance.

### Comparative analysis of viral RNA-sensing pathway reveals differences in basal and induced expression levels of key genes between bat and human

One suggested mechanism of greater disease resistance in bats is heightened innate immune response that is manifested in relatively high expression levels of type-I IFN or IFN-stimulated genes in bat cells and tissues^29, 31, 42, 81–83^. A recent study suggested that cells of Black Flying Fox (*Pteropus alecto*), a fruit bat from the Pteropodidae family (the same as *Rousettus*), show increased levels of the IRF transcription factors across bat tissues as well as higher basal expression of a set of IFN-stimulated genes^31^. We thus compared the expression of these genes across eight analogous tissues in *Rousettus*, human and mouse, using gene expression data from the Bat1K project^84^ (details on all samples used are available in **Table S17**), from GTEx^85^ and from the BodyMap transcriptomics dataset^86^, respectively, following data normalization to enable this comparison (see **Methods** for details on data processing and **Table S9** for detailed gene expression data). We observe no consistent trend in expression in this particular set of IRGs when comparing human and mouse with *Rousettus*, and a lower expression of some of the genes in *Rousettus*, suggesting divergence between *Pteropus* and *Rousettus* in these transcriptional patterns (**Supporting Figure 5**).

We hypothesized that other pathways relevant to viral infection may exhibit differential gene expression between human and bat, and focused on two major cellular pathways initiated by dsRNA-recognition: the RLR- and the TLR3-mediated pathways^87^. We observe that basal expression levels of some of the genes belonging to these pathways exhibit significant differences across tissues (**Fig 3A-C**). While some genes, such as IRF3 and LGP2 (an upstream regulator of two dsRNA sensors - MDA5 and RIG-I) are expressed in significantly higher levels in human tissues, other genes, including three major dsRNA sensors - TLR3, MDA5 and RIG-I are more highly expressed in bats (see also **Supporting Figure 6**, where human, mouse and *Rousettus* orthologous gene expression is shown). We note that we tested the transcriptional behavior of these genes in our single-cell data that includes human and *Rousettus* cells stimulated with dsRNA and profiled in parallel. These data appear in **Supporting Figure 7**, and while some gene differ in behavior between bulk and single-cell RNA-seq, as expected due to the different platforms and data used, they are in a significant agreement with each other (ρ=0.761 and P-value=2×10^-4^, and ρ=0.882 and P-value=2.8×10^-6^ between fold change values measured in bulk and single-cell RNA-seq in TLR- and RLR-associated genes, in human and *Rousettus*, respectively).

**Figure 3:**
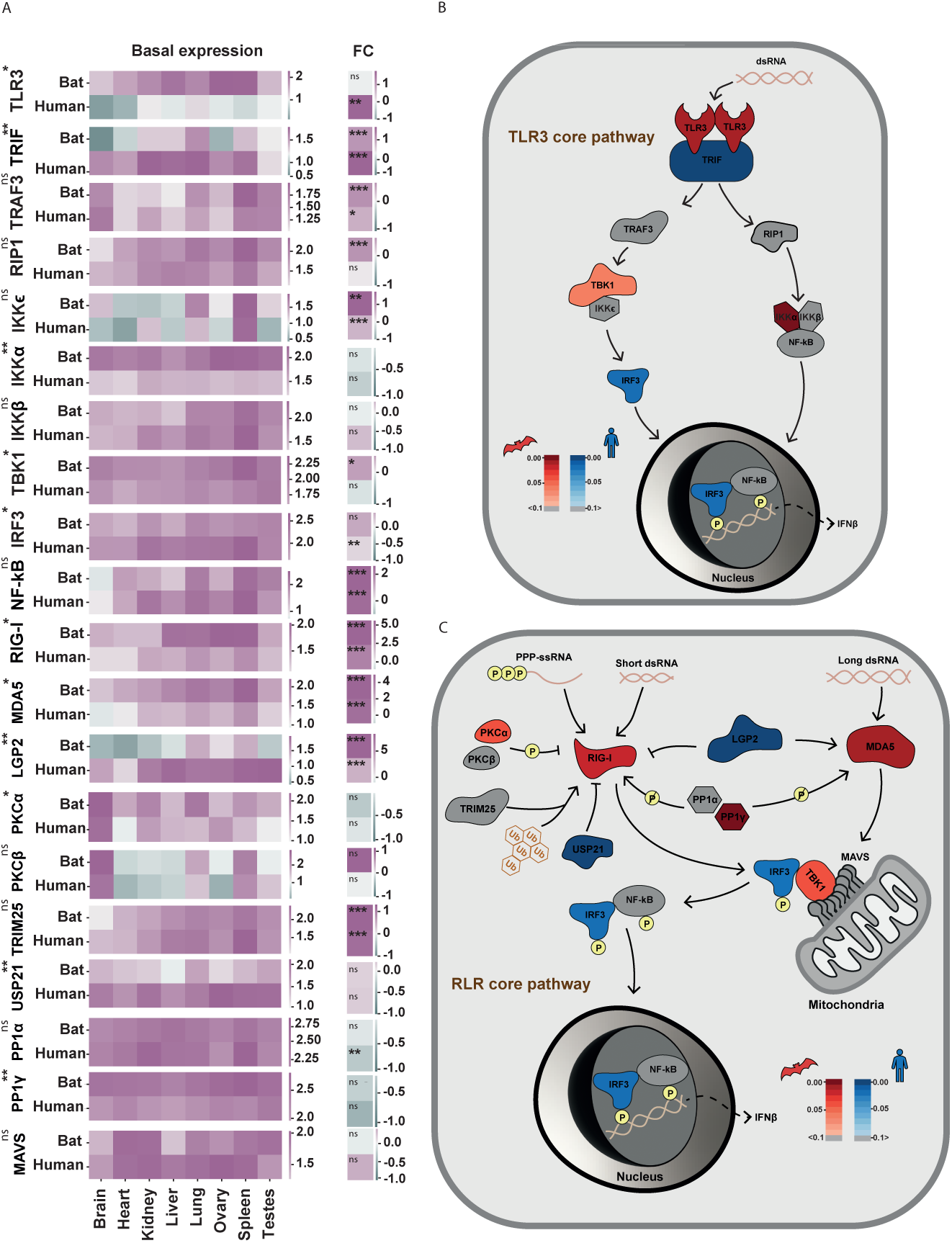
Basal expression and dsRNA-upregulation of genes in dsRNA-sensing pathways in *Rousettus* and human. (A) Normalized gene expression of orthologous genes in human and *Rousettus* across eight analogous tissues. Genes shown from the dsRNA-sensing core TLR3 and RLR pathways. Left: Normalized gene expression of orthologs in *Rousettus* and human in eight tissues. A one-sided paired T-test was used to test whether the ortholog genes are expressed more highly in one species (significance is shown to the left: ns = P>0.05, * = P<0.05, ** = P<0.01, *** = P<0.001). Right: log2(Fold Change) of the two orthologs in response to dsRNA, as measured in fibroblasts from human and *Rousettus* (significance of upregulation is shown within the square). **(B-C)** The TLR and RLR pathways, with genes that are significantly higher in basal expression across tissues in either human or *Rousettus* shown in red and blue, respectively (color scale denotes significance).

Finally, TLR3 displays a particularly interesting transcriptional behavior (**Figure 3A**): it is more lowly expressed in basal conditions in human tissues, but is strongly induced in response to dsRNA. In contrast, in *Rousettus*, TLR3 is more highly expressed in basal conditions across tissues, but is not significantly upregulated in dsRNA-stimulation, suggesting a switch between constitutive and induced expression between the two species.

### Bat-specific duplicates of innate immune genes display transcriptional divergence across tissues and rapid coding sequence evolution at the interface with viral proteins

The above analyses focused on one-to-one orthologs across species. Gene duplication was previously suggested to be an important mechanism in the evolution of the immune system^55^. We thus asked whether antiviral genes that rapidly diverge in transcriptional response to dsRNA (as measured by us between the two bat species) and that also display rapid coding sequence evolution (as shown in **Figure 2A-B**), belong to gene families that display relatively high duplication rates in the course of evolution. We observe that genes with a high level of transcriptional divergence do not display a significantly different duplication rate in comparison with low-divergence genes (**Supporting Figure 8**), suggesting that gene duplication is not significantly higher in the group of rapidly evolving bat antiviral genes as a whole.

We next searched for cases of bat-specific duplications or loss of innate immune genes, by scanning the list of dsRNA-upregulated genes in *Rousettus and Pipistrellus,* as well as those found to be upregulated in human and mouse. In addition to several antiviral genes that have been reported previously to be duplicated in bats, including the APOBEC3 family^84, 88^ and IFNω^42^, we also found that Tetherin (BST2) - an important restriction factor against a range of enveloped viruses has experienced rapid duplication in the branch that leads to *Pipistrellus* but not to *Rousettus*. In addition, we found that PLAAT4, a phospholipase A1/2 and an acyltransferase associated with cell proliferation and differentiation, that is significantly upregulated in response to dsRNA in *Pipistrellus*, has at least two copies in *Pipistrellus* (termed LOC118707299 and LOC118707212) - both of which are significantly upregulated in response to dsRNA. Related bat species - *Rhinolophus ferrumequinum* and *Myotis lucifugus -* are predicted to have 2 and 16 PLAAT4 duplicates in their genomes (as observed by profiling the orthologs and paralogs of these genes in ENSEMBL). In contrast, *Rousettus* and a related fruit bat, *Pteropus vampyrus* seem to have no copy of PLAAT4 in their genomes, suggesting rapid changes in copy numbers of this gene in different bat families. Similarly to this loss, both IFI44 and IFI44L genes, two IFN-stimulated genes associated with antiviral activities against several RNA viruses as well as with several autoimmune diseases^89–92^, are lost in *Rousettus* and in *Pteropus vampyrus* (the latter was observed in ENSEMBL genome annotation). We note that our analysis may be affected by genome quality, annotations and completeness, and we thus report only on cases of gene loss or duplication observed in more than a single bat genome.

We investigated the transcriptional and coding sequence divergence between gene duplicates of a specific gene - TNFRSF14, that has seven gene copies in *Rousettus (*LOC107499783,LOC107509685,LOC107510671, LOC107510672, LOC107510674, LOC107510998 and LOC107520019). In human cells, TNFRSF14 induces both pro- and anti-inflammatory pathways, based on binding to two different sites in its extracellular domain^93^. In addition, it serves as a receptor for entry of both human Herpes Simplex virus 1 and 2, through interaction with the herpes glycoprotein D (gD)^94^. TNFRSF14 is upregulated in response to dsRNA and to IFNB in human and mouse cells (**Figure 4A-B**). However, only one of its *Rousettus* gene copies is induced in response to dsRNA-stimulus. Interestingly, this copy is lowly expressed across tissues, in contrast to other *Rousettus* duplicates that display higher expression in two or more tissues. Furthermore, these paralogs seem to diverge in response to infection with Sendai and Marburg viruses (**Figure 4B, right**) (see a similar analysis on transcriptional divergence of APOBEC3 paralogs in **Supporting Figure 9**).

**Figure 4:**
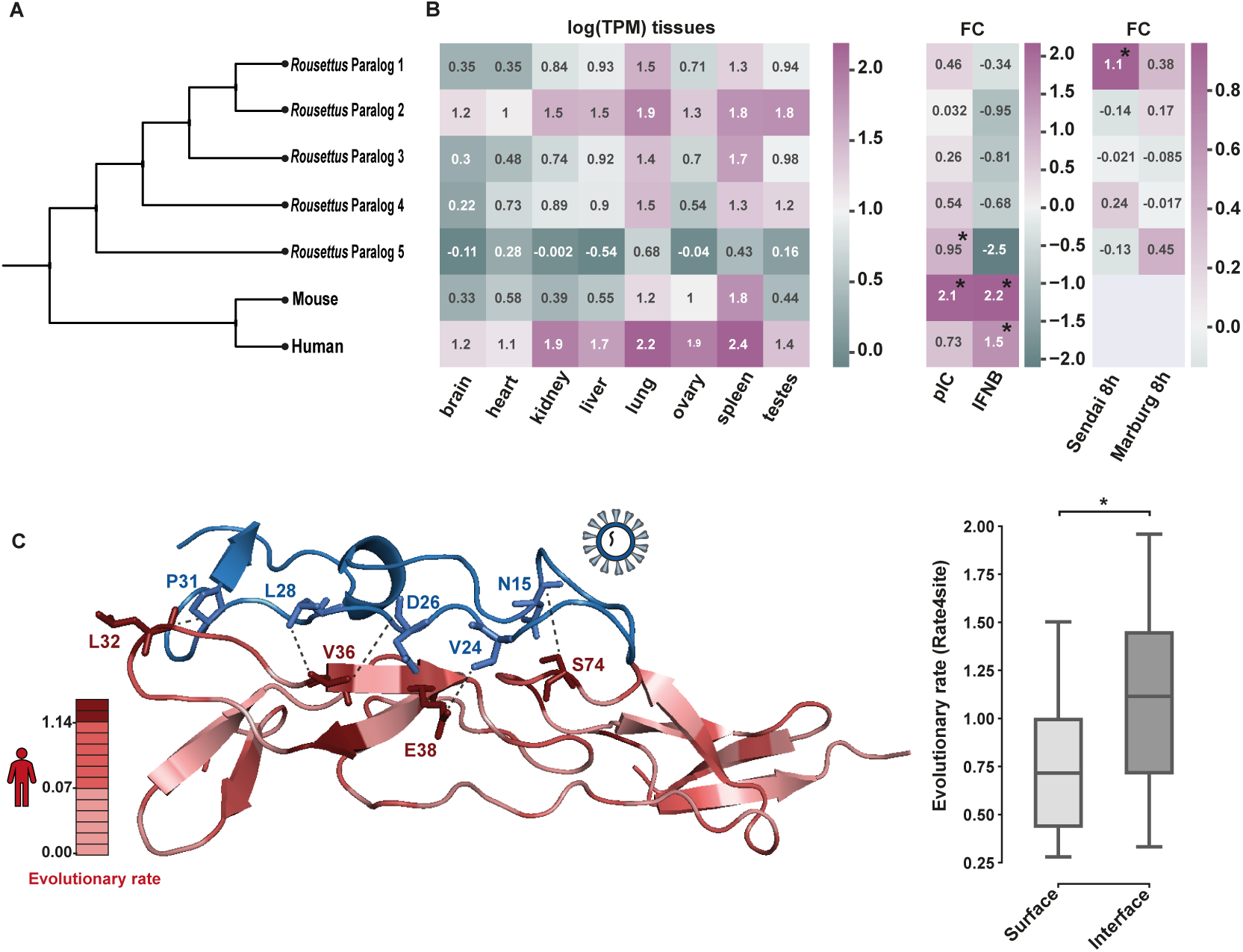
Duplication of TNFRSF14 in *Rousettus* and duplicate divergence in sequence and expression. (A) Reconstructed tree of TNFRSF14 of human, mouse and five *Rousettus* duplicates. **(B)** Left: TNFRSF14 gene expression across tissues of human, mouse and five *Rousettus* duplicates; Middle: log2(Fold Change) of TNFRSF14 of human, mouse and five *Rousettus* duplicates in response to dsRNA or IFNB, as measured in cells from human and *Rousettus* (significance of upregulation is shown within the square). Right: log2(Fold Change) of five *Rousettus* duplicates in response to infection by Sendai virus or Marburg virus. **(C)** Left: Structure of human TNFRSF14 with the herpes glycoprotein D, in red and blue respectively (PDB: 1JMA). Human TNFRSF14 is colored by relative evolutionary rate of residues (as measure across all orthologs and paralogs appearing in A). Right: The distributions of evolutionary rates of residues found at the surface of TNFRSF14 (surface) compared with evolutionary rates of residues interacting with the herpes glycoprotein D (interface). Comparison is made using a Mann-Whitney test (* = P<0.05).

In addition to the observed transcriptional divergence between the *Rousettus* TNFRSF14 duplicates, we investigated the divergence in coding sequence between these duplicates. We estimated the relative divergence in amino acid substitution across the *Rousettus* TNFRSF14 duplicates in each amino acid position, using Rate4site^95^ (see **Figure 4C** and Methods). We then contrasted the substitution rates of residues known to be part of the interface formed between human TNFRSF14 and herpes glycoprotein D proteins, and the other surface residues (identifying the interface residues using the CSU method^96^, as we done previously^97^). We observed a significantly higher rate of substitution in the interface residues in comparison with the surface residues (**Figure 4C**). This suggests that this region, known at least in human to be a site of interactions with viral proteins, diverged more rapidly in sequence between *Rousettus* paralogs. This analysis demonstrates the potential of recent gene duplicates to diverge in both sequence and expression, and potentially to contribute to bat-specific response against viruses.

## Discussion

Some of the unique physiological and behavioral characteristics of bats, are likely linked to their co-evolution and adaptation to viral infections. Here, we took a comparative transcriptomics approach to study the expression program triggered as part of the immediate antiviral response in bats and the evolution of specific antiviral genes. We profiled this innate immune response in fibroblast cells from two bat species - Kuhl’s pipistrelle (*Pipistrellus kuhlii*) and the Egyptian fruit bat (*Rousettus aegyptiacus*) - and compared this response between these two bat species as well as between bats and other mammals.

We observe that the overall innate immune response is conserved across bats, where genes are up- and downregulated in response to dsRNA in a similar manner to other mammals both with respect to the overall numbers of regulated genes and in terms of specific gene identities and cellular pathways involved in this response. When comparing the level of response across species, by looking at gene’s fold change in response to dsRNA in pairs of one-to-one orthologs, we observe that the response of the two bat species is conserved, showing a significant correlation between ortholog responses in the two species. However, this level of correlation is similar to the correlation observed between primates and rodents, and lower to than correlations within the rodent or within the primate clade – exemplifying the divergence between distant bat species in the level of upregulation of specific genes. These results also emphasize the importance of further studies into separate branches of the bat clade that likely evolved differently in their diverse habitants in response to different viral strains.

Our focused analysis of genes that are transcriptionally divergent in their response to dsRNA between the two bat species revealed several regulatory and evolutionary trends: genes diverging in transcriptional response between the two bat species tend to also rapidly diverge in coding sequence. This suggests that innate immune genes in bats evolve concurrently through distinct evolutionary mechanisms, including changes in the level of expression and in amino acid substitutions across species. These findings are in line with our previous observations where we studied the evolution of antiviral genes in primates and rodents and showed that these transcriptionally diverging antiviral genes also diverge rapidly in coding sequence^56^. When looking at the promoter architecture of innate immune genes in *Rousettus*, we observe that in agreement with previous findings in various mammalian systems^77, 78, 98^, gene transcriptional divergence between species is associated with particular promoter structures: Genes with CGIs in their promoters tend to be transcriptionally conserved between species whereas genes without such elements, and especially those with a TATA-box in their promoters, tend to be divergent in their transcriptional response between the two bat species. Furthermore, transcriptionally divergent bat genes also show high cell-to-cell variability in expression between individual cells within the same species. This is true for both *Rousettus* and human cells, and in both stimulated and unstimulated cells, as revealed by our single-cell transcriptomics of human and bat cells. The similarity observed between human and *Rousettus* cells in the set of genes found to be transcriptionally divergent (i.e., genes found to be divergent in bat cells are also divergent in human cells), suggests that the same set of innate immune genes have undergone rapid transcriptional evolution in the course of mammalian evolution while other genes involved in this response are conserved. This is likely due to different levels of regulatory constraints imposed on these genes, such as their function (whether they have pleotropic or regulatory functions, such as transcription factors that are more conserved^56^) and the biophysical nature of their protein products – whether they are engaged in direct interactions with viral proteins and what constraints limit the evolution of the residues that interact with the viral proteins^97^.

Another difference between species that may give rise to differences in their ability to resist pathogen infection is the basal expression of innate immune genes. This mechanism of greater resistance has been suggested in *Pteropus alecto*. In *Rousettus*, a previous analysis suggested greater tolerance to infection by Marburgvirus^15^, a virus for which *Rousettus* is the natural reservoir and that leads to mild subclinical symptoms in *Rousettus* upon infection. In this work, it was shown that while an antiviral program is triggered upon infection, several important inflammatory genes including CXCL8 and IL6 are not induced, suggesting a specific upregulation of defense programs without eliciting a strong inflammatory response that can lead to tissue damage^15^. Here, we compared basal expression of genes involved in dsRNA sensing, including viral sensors from the RLR and TLR families and their downstream effectors, in *Rousettus,* human and mouse. We observe that several of these genes are more highly expressed in bat tissues, whereas other genes in these pathways display an opposite trend. Together with their known adaptative coding sequence evolution (in all RLRs, and is some TLRs^99–103)^, this divergence in basal expression suggests that genes involved in viral sensing pathways in bats evolved through altered expression levels in addition to changes in coding sequences across species, potentially contribute to bat antiviral immunity.

In contrast to previous findings based on human and non-human primates^55, 56^, we do not observe that the most transcriptionally divergent innate immune genes in our data belong to gene families that have undergone rapid duplication. This is in agreement with the relatively few genes known to be involved in antiviral immunity that were found to rapidly duplicate or to have been lost in the bat clade in comparison with other genes^84^.

Focusing on specific genes that are upregulated in response to dsRNA and that have relatively high copy numbers in bats in comparison with other mammals, we found several genes and gene families previously not reported to undergo rapid gene duplication or gene loss in bats. These include both IFI44 and IFI44L genes that seem to have been lost in Pteropodidae, and PLAAT4 that shows a wide variation in paralog numbers in Vespertilionidae. We further analyzed the transcriptional and coding sequence divergence of an additional innate immune gene, TNFRSF14, that has undergone rapid duplication in the branch leading to *Rousettus*. The paralogs of this gene in *Rousettus* display distinct patterns of basal expression across bat tissues, as well as in their response to IFN and to dsRNA. Furthermore, when these paralogs were compared with the single orthologs found in human and mouse genome, we observed that the relatively high basal expression and upregulation of the single copy of TNFRSF14 in human and mouse differ from what observed in TNFRSF14 bat paralogs, suggesting transcriptional divergence between paralogs. When looking at the substitution rate of amino acids across paralogs of bat TNFRSF14, we observe an interesting trend: The residues whose homologs in human TNFRSF14 were shown to form an interface with herpes glycoprotein D (gD) (based on the crystal structure of the human-herpes virus complex), are significantly more divergent in comparison with other surface residues of the TNFRSF14 protein. These analyses demonstrate how recent duplication events of various bat immune genes in specific branches of the bat clade, rapidly diverged through basal expression level across tissues, in transcriptional response to infection, and in coding sequence, in regions that may be involved in direct contact with viral proteins. This example, as well as recent work on functional diversification of PKR duplicates in *Myotis* species^38^ and of BST2 (Tetherin) duplicates in *Vespertilionidae*^39^, suggest that rapid duplication of specific antiviral genes has contributed to bat-specific adaptation against infection from specific viral strains or viral families.

Our work provides a comparative catalog of antiviral gene expression using a large set of individuals in two bat species. These data give insights into how different evolutionary mechanisms, including changes in gene expression in homeostasis and during infection, as well as coding sequence evolution and gene duplication, may have shaped the innate immune response of bats and contributed to bat adaptation to viral infections. In addition to providing an important resource for the community interested in bat immunity, these data have the potential to advance our understanding of pathological immune conditions in humans.

## Methods

### Ethical compliance

Bat biopsies from two bat species - *Rousettus aegyptiacus* and *Pipistrellus kuhlii* - were collected from either naturally deceased individuals or from animals scarified as part of a different project, with approvals by the Israel Nature and Parks Authority (approval number 2020/4285) and the TAU ethics committee (approval number 04-20-023).

### Bat dermal fibroblast extraction and growth

Bat dermal fibroblast cells were extracted from skin samples and processed in a similar manner to our previously described protocol^56^. Following extraction, we passaged cells for three passages, using a growth medium DMEM (Biological industries, 01-052-1A) supplemented with 20% FBS (Rhenium, 10270106) 2 mM L-Alanyl-L-Glutamine (Biological industries, 03-022-1B), 1 mM Sodium Pyruvate (Biological industries, 03-042-1B), Primocin (Invivogen, ant-pm-1) and Penicillin-Streptomycin-Amphotericin-B (Biological industries, 1 03-033-1C. Cells were then frozen and stored in liquid nitrogen. Individual lines that did not grow or showed signs of senescence were discarded. In the case of *Rousettus*, skin biopsies from shoulders were also collected and grown in a similar fashion. See full list of samples in **Supporting Table 1**. For single-cell experiments, we also used a primary cell line of human dermal fibroblast from ATTC (PCS-201-012).

### Bat cell stimulation with dsRNA

Prior to stimulation, cells were thawed and grown for a few days in fibroblast growth medium. A day before stimulation, cells were trypsinized, counted and seeded into 6-well plates to reach ∼ 70% confluence at the start of stimulation. Cells were stimulated as follows: (1) stimulated with 1 μg/ml high-molecular mass poly(I:C) (Invivogen, tlrl-pic) transfected with 2 μg/ml Lipofectamin 2,000 (ThermoFisher, 11668027); (2) mock transfected with Lipofectamin 2,000. After 4 hours, the stimulation was terminated by washing the cells in PBS. For bulk RNA-seq, cells were lysed using RNA Lysis Buffer (Zymo Research). For droplet-based single-cell RNA-seq, cells were first trypsinized and then collected to continue with library preparation according to the manufacturer’s protocol for the Chromium Single Cell 3′ gene expression v.3.1 (10x Genomics), as described below.

### Bulk RNA-seq

Total RNA was extracted using the Quick-RNA MicroPrep kit (Zymo Research, Cat. No. R1051). RNA concentration, purity and integrity were measured with Qubit 3.0 and TapeStation 4200. In total, 49 samples were used for subsequent library preparation and sequencing.

Libraries were produced using the Kapa Stranded mRNA-seq Kit (24RXN) (KK8420 ROCHE-07962193001) with KAPA Unique Dual-Indexed Adapter Plate (KK8726-08861862001) and KAPA Adapter Dilution Buffer (KK8721-08278539001), starting with up to 500ng of total RNA. Final cDNA amplification was performed with 13 PCR cycles. Libraries were normalized and pooled at 10nM. Pooled samples were sequenced on an Illumina NovaSeq 6000 instrument, using the NovaSeq 6000 SP Reagent Kit (200 cycles) 20040326 and paired-end 101-bp reads.

### Single-Cell RNA-seq

Following trypsinization and cell count, human and Rousettus samples of the same treatment condition (poly(I:C) and lipofectamine) were pooled together. Cells were prepared and loaded to the Chromium controller according to the manufacturer’s protocol (10x Genomics), using the Single Cell 3’ Reagent Kit (version 3.1). cDNA synthesis and amplification were performed according to the protocol. Libraries were sequenced on an Illumina NovaSeq 6000 using the NovaSeq 6000 SP Reagent Kit v1.5 (100 cycles).

### Read mapping to annotated transcriptome

#### Bulk RNA-seq

Reads were mapped and gene expression was quantified using Salmon^104^ (version 1.4.0) with the following command parameters for each sample: ‘salmon quant -i [index_file_directory] -l IU −1 [left_read_library/lane1] [left_read_library/lane2] −2 [right_read_library/lane1] [right_read_library/lane2] --geneMap [transcript_to_gene_file] --seqBias --gcBias -q --numBootstrap 100 --threads 8 --validateMappings -o [output_directory]’. Each sample was mapped to its respective species’ annotated transcriptome, downloaded from the NCBI website (*Rousettus aegyptiacus*: Genome published by Pavlovich et al.^36^; *Pipistrellus kuhlii*: from the Bat1K project^84^). Following quantification, a count matrix was created.

In the case of *Rousettus aegyptiacus*, we mapped reads to two genome annotations – the genome published by Pavlovich et al., and the genome from the Bat1K project. Since we had higher mapping rates of reads using the first genome, we based our subsequent analyses on that mapping.

We also mapped previously published datasets of bulk RNA-seq of *Rousettus aegyptiacus* samples: (1) IFN stimulation of *Rousettus* cell lines from Pavlovich et al. (2020)^42^ (run accessions SRR11148658-723), (2) Viral infection of *Rousettus* cell lines from Arnold et al. (2018)^10^ (run accessions SRR7548028-63) and (3) *Rousettus* tissues dataset from the Bat1K^84^ were downloaded from the NCBI Sequence Read Archive and mapped to the Rousettus UCSC transcriptome. Paired read libraries were mapped as above, but with libType A. Single read libraries were mapped as follows: ‘salmon quant -i [index_file_directory] -l A -r [sample_library] --geneMap [transcript_to_gene_file] --seqBias --gcBias -q --numBootstrap 100 --threads 8 --validateMappings -o [output_directory]’.

RNA-seq data of tissues from human and mouse was used in comparison with *Rousettus* tissue gene expression. These data were downloaded from GTEx^85^ and BodyMap^86^ for human and mouse, respectively, and processed as previously described^105^. To achieve comparable gene expression data across species, we filtered all the pseudo autosomal expression records and merged various brain tissues by computing the mean for each gene across them, to compare with the mouse and bat brain data. For each gene in each of the eight tissues that existed in all three species, we used the mean expression between all male and female individuals as representative value of this species.

#### Single-Cell RNA-seq

Reads were demultiplexed and quantified using Cell Ranger from Chromium Single Cell Software Suite (version 5.0.1, 10x Genomics Inc). Raw read data was demultiplexed (“unpooled”) into sample-specific FASTQ files based on the sample indices. Reads were then quantified with the following command for each sample: ‘cellranger count --id=[output_directory] --transcriptome=[reference_genome_directory] --fastqs=[sample_FASTQs_directory]’. Each sample was mapped to its respective species’ annotated genome, the *Rousettus aegyptiacus* genome (see above) and the human GRCh38 reference genome (ENSEMBL version 99), as well as to an index that included of both genomes (human and Rousettus). The latter was initially used to determine whether each individual cell originated in human or Rousettus. Cells were determined to be from *Rousettus*, if more than 80% of their reads were mapped to Rousettus, and human cells were determined in a similar fashion. Cells that did not meet these criteria were considered doublets and were discarded. Following species determination for each single-cell library, all subsequent analyses were based on the separate mapping output.

### Quantifying differential gene expression in response to dsRNA

#### Bulk RNA-seq

We quantified differential gene expression between dsRNA-treatment and control (for each species separately and, in the case of *Rousettus*, for each skin tissue separately as well) by using edgeR (version 3.32.1)^65^ with rounded estimated counts from Salmon. We only kept genes that were expressed in at least 3 of the 30 *Rousettus* samples or 2 of the 19 *Pipistrellus* samples. In each of the public datasets (IFN stimulation, viral infection), we kept genes expressed in at least 3 of the respective dataset’s samples. Differential expression analysis (DEA) was conducted via the edgeR exact test. P-values were corrected for multiple testing by estimating the false discovery rate (FDR). For the public datasets, separate DEAs were performed for each time point and each virus or IFN type.

#### Single-Cell RNA-seq

We filtered cells with less than 10,000 reads, and genes expressed in less than 3 of the remaining cells. We then removed cells with more than 20% mitochondrial reads. Further single-cell data analysis was done using Seurat (version 4.0.1)^106^. Cell cycle phase was regressed out via Harmony^107^. Following lowly expressed gene filtering, 75% of the most variable genes were used for dimensionality reduction. Differentially expressed genes in stimulated versus unstimulated cells were estimated using Seurat’s FindMarkers function.

### Orthology mapping between bats, primates and rodents

To determine gene orthology relationship between species as accurately as possible, we integrated orthology annotations from ENSEMBL Compara^108^ for human, rhesus macaque, mouse and rat genes, while finding orthologs between human and the two bat species using eggNOG (version 2.1.2)^72^.

To obtain gene orthology using eggNOG, we ran the following command for human and mouse genomes (ENSEMBL version 99, after removal of secondary haplotypes) as well as the CDS files of the two bat species (with the genome versions mentioned above): ‘emapper.py --data_dir {database_dir} --cpu 6 -i {input_cds_file} --itype CDS --output {output_name_for_results} --sensmode very-sensitive --report_orthologs --output_dir {results_dir} -m diamond -d none --tax_scope 40674 --go_evidence non-electronic --target_orthologs all --seed_ortholog_evalue 0.001 --seed_ortholog_score 60 --query_cover 20 --subject_cover 0 --override --scratch_dir {scratch dir} --temp_dir {tmp dir}’.

Genes were defined as belonging to the same orthology group (either one or more from each species), if their transcripts mapped to the same eggNOG name. Genes whose transcripts were mapped to more than a single eggNOG name were removed. The detailed eggNOG processed results for human, mouse, *Pipstrellus* and *Rousettus* (two genome annotations) are in **Table S8**.

From these cross-species mappings we created several tables of one-to-one orthologs – in Table S9: For cross-mammalian analysis (involving six species, as in **Figures 1-2**), we used only genes that had one-to-one orthology annotations in ENSEMBL for the primates and rodents, as well as one-to-one orthology mappings in eggNOG for human, *Rousettus* and *Pipistrellus* (10,241 genes). For cross-bat analysis we used one-to-one orthology annotations between the two bat species (13,888 genes).

### Fold-change-based analysis of conservation and divergence in innate immune response

To compare the overall change in response to immune stimulation between pairs of species, we computed the Spearman’s rank correlation of the fold-change between all one-to-one orthologs that were differentially expressed in response to dsRNA in at least one species (q-value<0.01 in one of the 6 species-2,865 genes) as well as of all expressed genes (**Figure 1E-H**).

### Quantifying transcriptional divergence in innate immune response between species

To estimate transcriptional divergence between the six species in response to treatment, we focused on genes that have one-to-one orthologues in all tested species. We defined a measure of response divergence between the two bat species or between bat and human, by calculating the absolute differences in the fold-change estimates across the two orthologues, for genes that are differentially expressed in at least one species, as previously done^56, 109^. For example, between the two bat species, we defined: *Response diveregnce* = |*FC_Rousettus_* - *FC_Pipistrellus_*|. We classified the top third most divergent genes as high divergent genes, the medium third as medium divergent genes, and the bottom third as low divergent genes. In several analyses, the top 20% genes highly responding in *Rousettus* or *Pipistrellus* with respect to the other bat species were used (in the section: **Characteristics of *Rousettus*-specific and *Pipistrellus*-specific antiviral genes**). In these analyses we used the same Response divergence measure only without using the absolute value to determine which genes are species-specific.

### Promoter sequence analysis

Human and *Rousettus* CGI annotations were downloaded from the UCSC genome table browser^110^ (hg38 for human), and CGI genes were defined as those with a CGI overlapping their core promoter (300bp upstream of the TSS reference position and 100bp downstream of it, as suggested previously^56, 77^). Genes were defined as having a TATA box if they had a significant match to the Jaspar TATA box matrix (MA0108.1)^111^ in the 100bp upstream of their TSS by FIMO^112^ with default settings (we used a 100 bp window owing to possible inaccuracies in TSS annotations).

### Coding sequence evolution analysis

The dN/dS ratio (non-synonymous to synonymous codon substitutions) values of *Rousettus* genes across bats and their statistical significance were obtained from a previous study that used orthologous genes from 18 bat species (Hawkins et al., 2019)^76^. We computed distributions of the test statistics of positive selection, based on dN/dS values for each of the three groups of genes with low-, medium- and high-divergence in response to dsRNA (**Figure 2A**).

### Sequence similarity analysis

To obtain the level of similarity between orthologs, we ran BLAST of *Rousettus* versus *Pipistrellus.* We selected the longest CDS for each pair of one-to-one orthologs (and the lowest TSL, in case two transcripts had the same length) of the two species. We then ran blast2seq on these the CDS sequences to obtain the similarity level.

### Rate of gene gain and loss analysis

The significance (P-value) of a gene family to have undergone a high rate of gene duplication and loss (contraction) over the course of vertebrate evolution, in comparison to other gene families, was obtained from ENSEMBL^108^. These statistics are based on the CAFE tool that estimates the global birth and death rate of gene families and identifies gene families that have accelerated rates of gain and loss^113^. We calculated the distributions of P-values for the low-, medium- and high-divergence gene groups (as defined above), and plotted them as the negative logarithm values (**Supporting Figure S8**).

### Tolerance and Resistance analysis

Genes primarily associated with tolerance and resistance expression programs were taken from a previous study, which compared the transcriptional response and physiological outcomes of infection across a large set of mouse strains^80^. The association of the gene sets identified in that work with tolerance and resistance to infection were suggested to be conserved across mammals and infectious diseases, as demonstrated by further analyses of additional systems.

We used the top 20% *Rousettus-* or *Pipistrellus*-specific genes separately, with two measures of resistance and tolerance from the above-mentioned paper, i.e., coordinates on axis T+ R (**Figure 2F**) and Gene-to-T or R correlation (**Supporting Figure 4**). The detailed T and R data appears in **Supporting Table S16.**

### Cell-to-cell variability analysis

To quantify the biological cell-to-cell variability of genes, we used the DM (Distance to Median) approach — an established method that estimates relative cell-to-cell variability in gene expression while accounting for confounding factors such as gene expression level^79^ and after removal of lowly expressed genes, as detailed in a previous study^56^.

### Cross-species ortholog gene expression analysis across tissues

In order to compare one-to-one orthologous gene expression across analogous tissues between *Rousettus*, human and mouse (such as when looking at RLR- and TLR-related genes), we obtained gene expression of the three species in eight tissues as described above. We re-computed the values of ortholog gene expression as follows: we took all genes with one-to-one orthology relationship between human, mouse and bat (10,230 genes), and for each library we normalized the gene expression value by dividing it by the total gene expression across all orthologous genes (total TPM), multiplied by one million. We compared expression of these normalized values across tissues between pairs of species (*Rousettus* versus human, *Rousettus* versus mouse), by using a one-sided paired t-test across a vector of gene expression values in analogous tissues (using the ttest_rel function from scipy) and correcting by FDR.

### TNFRSF14 evolutionary analysis

The relative evolutionary rate of each amino acid in the TNFRSF14 protein was estimated using a multi-sequence alignment and phylogenetic tree using Rate4site^95^, as described previously^97, 98^. Briefly, five *Rousettus* paralog sequences and other species ortholog sequences (human, mouse and *Pipistrellus*) were aligned using MUSCLE^114^. A phylogenetic tree was constructed using PhyML^115^, and Rate4site was run using default parameters.

Interface and surface residues were inferred from the solved structure of TNFRSF14 with the herpes glycoprotein D (pdb: 1JMA^116^) using the CSU method^96^.

### GO term analysis

To study the functional enrichment of genes against the background of the entire human genome we used g:Profiler program^66^. In the section of ‘Characteristics of *Rousettus*-specific and *Pipistrellus*-specific antiviral genes’ we used GOrilla^117^ where we tried to study the functional enrichment of *Rousettus*-specific genes against the background of *Pipistrellus*-specific genes and vice versa. The detailed results of both g:Profiler (g:GOST-fuctional profiling and g:Convert-gene ID conversion) and GOrilla programs are in Supporting Tables S10-15.

### Single-cell pseudotime trajectory analysis

Following dimensionality reduction by Harmony^107^, the first dimension was taken as a pseudotime and selected gene trajectories are shown in **Supporting Figure 7**.

### Data availability

Sequencing data produced in this study are available in ArrayExpress with the following accessions: E-MTAB-10870 and E-MTAB-11056.

Major scripts can be found in: https://github.com/lilachschn/bat_gene_expression_divergence

## Acknowledgments

We would like to thank Xi Chen, Natalia Kunowska and Naama Peshes-Yaloz for helpful discussions, Hila Kobo and Rami Khosravi for technical assistance with NGS library preparations. This research was supported by the Israel Science Foundation (ISF, grant No. 435/20), the Chan Zuckerberg Initiative (Single-Cell Analysis of Inflammation, grant No. DAF2020-217532), by a grant from the Zimin Institute for Engineering Solutions Advancing Better Lives, and by a joint QBI/UCSF-TAU research grant in computational biology and drug discovery. SK was supported by a Minerva Fellowship of the Minerva Stiftung Gesellschaft fuer die Forschung mbH.

