## Supporting Information for "A comparative analysis of the antiviral response in two bat species reveals conserved and divergent innate immune pathways"

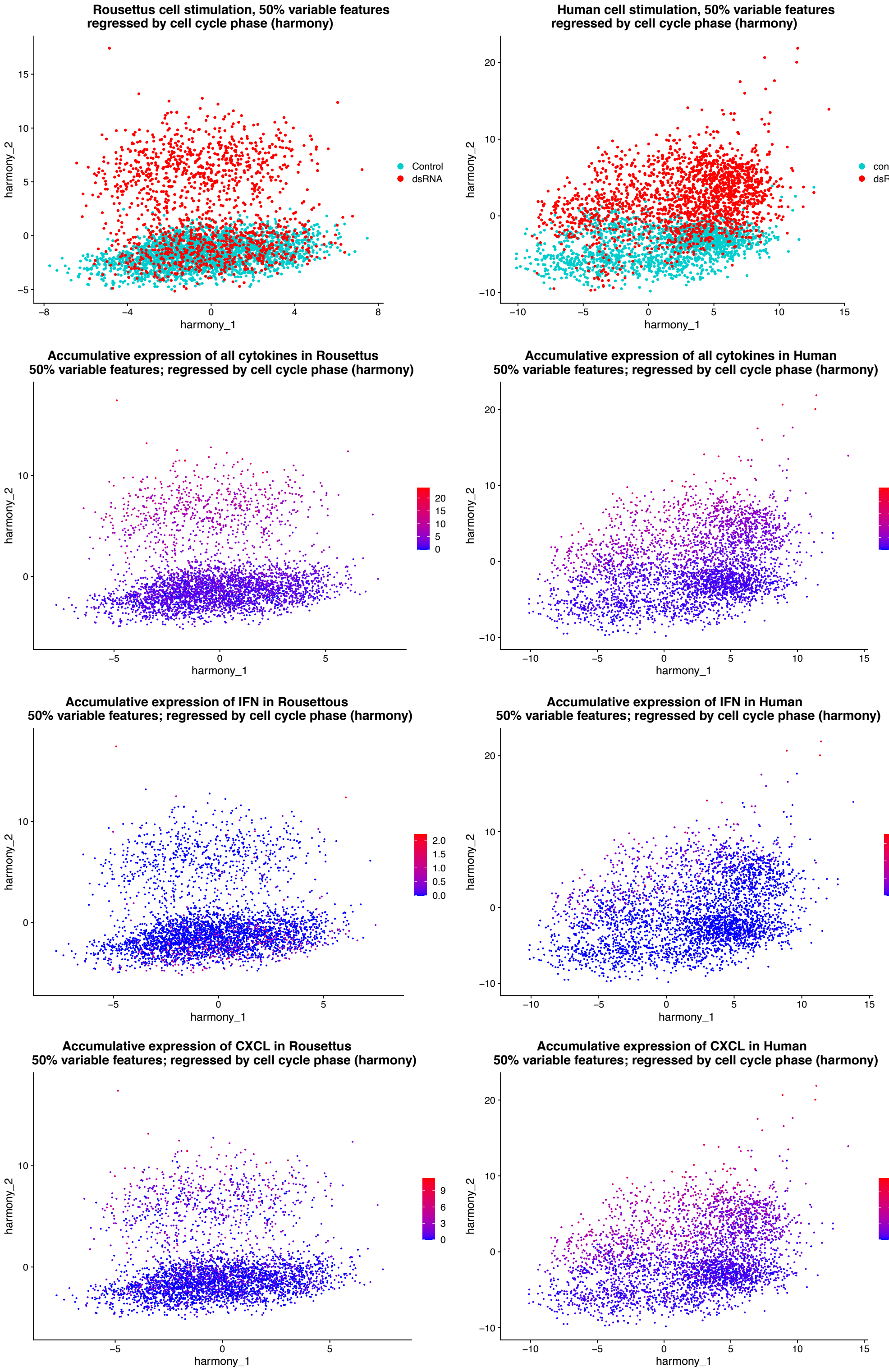


**Supporting Figure 1: Cytokine expression in single-cell.** Cells are shown following dimensionality reduction using Harmony. Expression of all cytokines, only interferons (IFN) or only CXCL chemokines are shown for human and *Rousettus*. In the top panels the conditions of cells are shown – dsRNA (poly IC treatment) or Control (mock lipofectamine transfection).

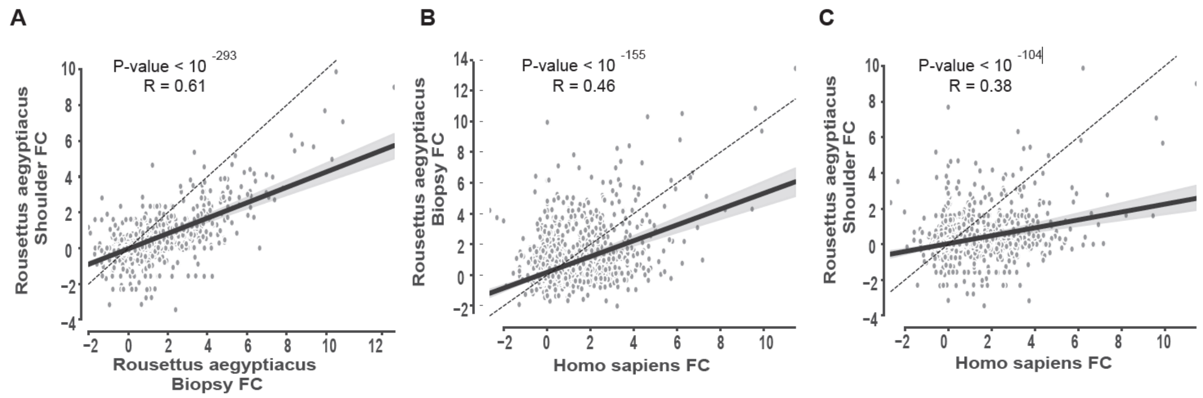


**Supporting Figure 2: Correlation of gene expression fold-change in response to dsRNA in different skin regions of *Rousettus* and between *Rousettus* and human. (A)** Spearman's rank correlation of 2,865 genes from *Rousettus* dermal fibroblasts extracted from skin from shoulder and from wing biopsy. **(B)** Spearman's rank correlation of 2,865 one-to-one orthologous genes between human fibroblasts and *Rousettus* dermal fibroblasts extracted from wing biopsies. **(C)** Spearman's rank correlation of 2,865 one-to-one orthologous genes between human fibroblasts and *Rousettus* dermal fibroblasts extracted from shoulder skin. In all cases, the set of genes includes genes that are found to be differentially regulated in response to dsRNA in at least one of the six studied species (q-value<0.01), as in Fig 1.

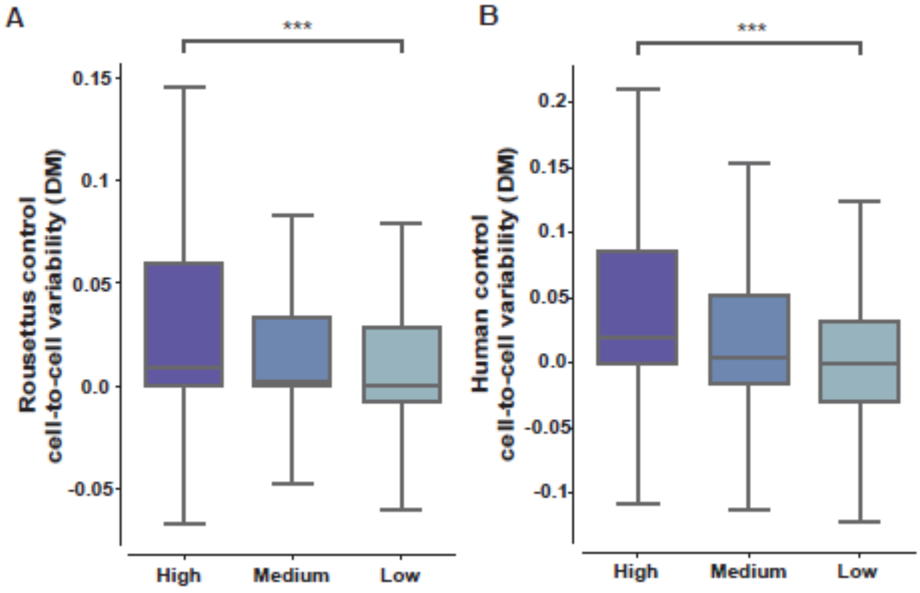


**Supporting Figure 3: Relationship between gene expression cell-to-cell variability and gene response divergence between species.** Genes are partitioned into three groups of low-, medium- and high-divergence in response to dsRNA between species, as explained in the main text. The distribution of cell-to-cell variability in gene expression, as measured using the distance to median approach (DM), is shown for each of these three groups for **(A)** Unstimulated *Rousettus* cells (2,030 genes with DM data) and **(B)** Unstimulated human cells (2,528 genes with DM data). High- and low-divergence groups are compared using a Mann-Whitney test (*** - P<0.001).

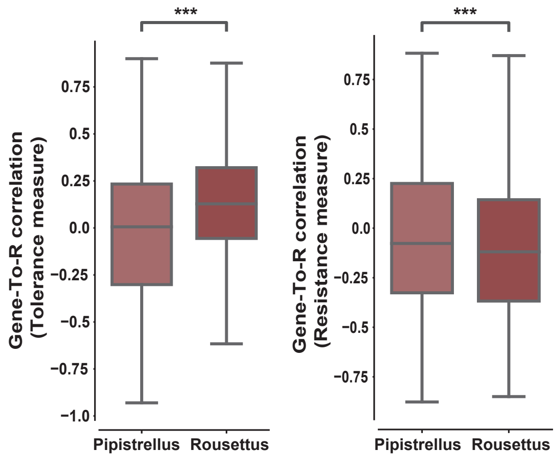


**Supporting Figure 4: Relationship between gene programs associated with tolerance and resistance and dsRNA-responsive genes that are highly responding in *Pipistrellus* and *Rousettus*.** Distribution of Gene T-To-R correlation axis of 1,949 *Pipistrellus*- and 1,928 *Rousettus*-specific genes, with the measures taken from a previous study(Cohn et al. 2022), and as explained in the main text. The distributions are compared using a Mann-Whitney test (*** - P<0.001).

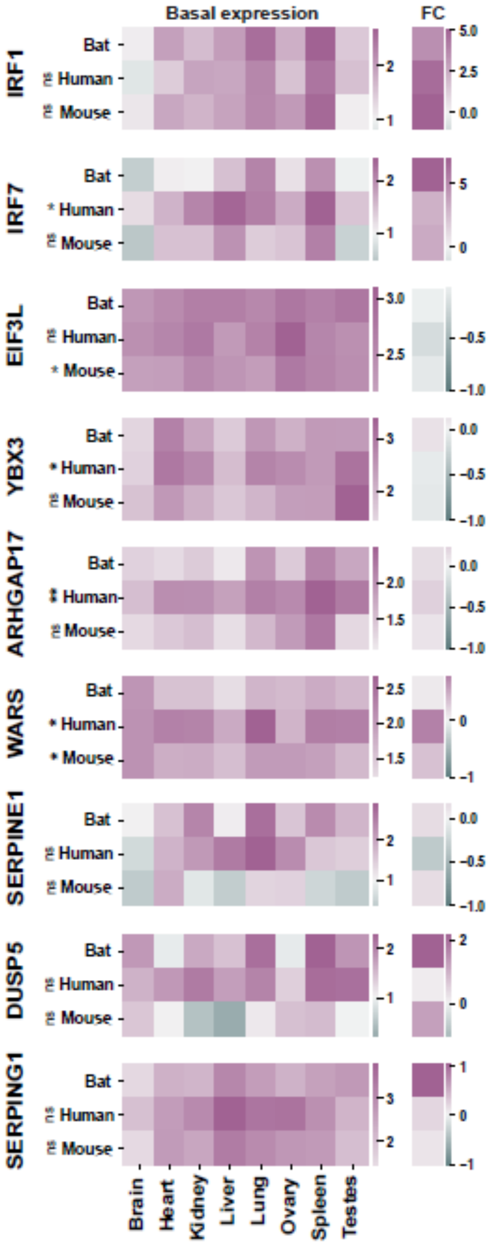


**Supporting Figure 5: Expression levels of orthologous genes previously shown to be highly expressed in Black Flying Fox, across eight tissues in human, mouse and *Rousettus*.** Genes previously reported (Irving et al. 2020) to be basally expressed in higher levels in Black Flying Fox, were tested for their basal expression in eight equivalent tissues in human, mouse and *Rousettus*. A comparison of their level of expression between Rousettus and human, and separately, between *Rousettus* and mouse was preformed using a one-side paired T-test, following gene expression normalization. FDR-corrected P-values are shown next to the human and the mouse row, respectively. (ns = not significant (P-value>0.05), * = P<0.05, ** = P<0.01). The separated right column shows each of these genes' fold-change in response to dsRNA, as measured in dermal fibroblasts.

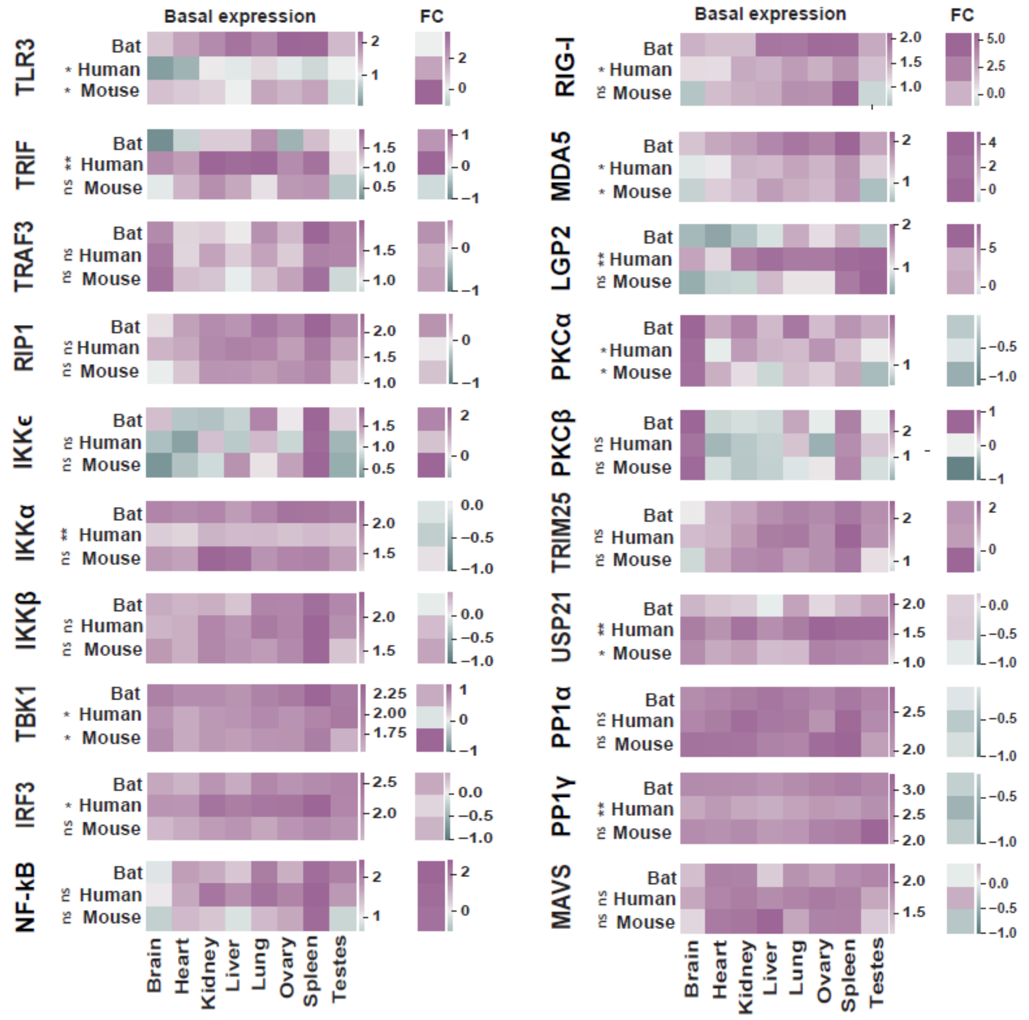


**Supporting Figure 6: Expression levels of orthologous genes in the RLR and TLR pathways, across eight tissues in human, mouse and *Rousettus*.** Genes were tested for their basal expression in eight equivalent tissues in human, mouse and *Rousettus*. A comparison of their level of expression between Rousettus and human, and separately, between *Rousettus* and mouse was preformed using a one-side paired T-test, following gene expression normalization. FDR-corrected P-values are shown next to the human and the mouse row, respectively. (ns = not significant (P-value>0.05), * = P<0.05, ** = P<0.01). The separated right column shows each of these genes' fold-change in response to dsRNA, as measured in dermal fibroblasts.

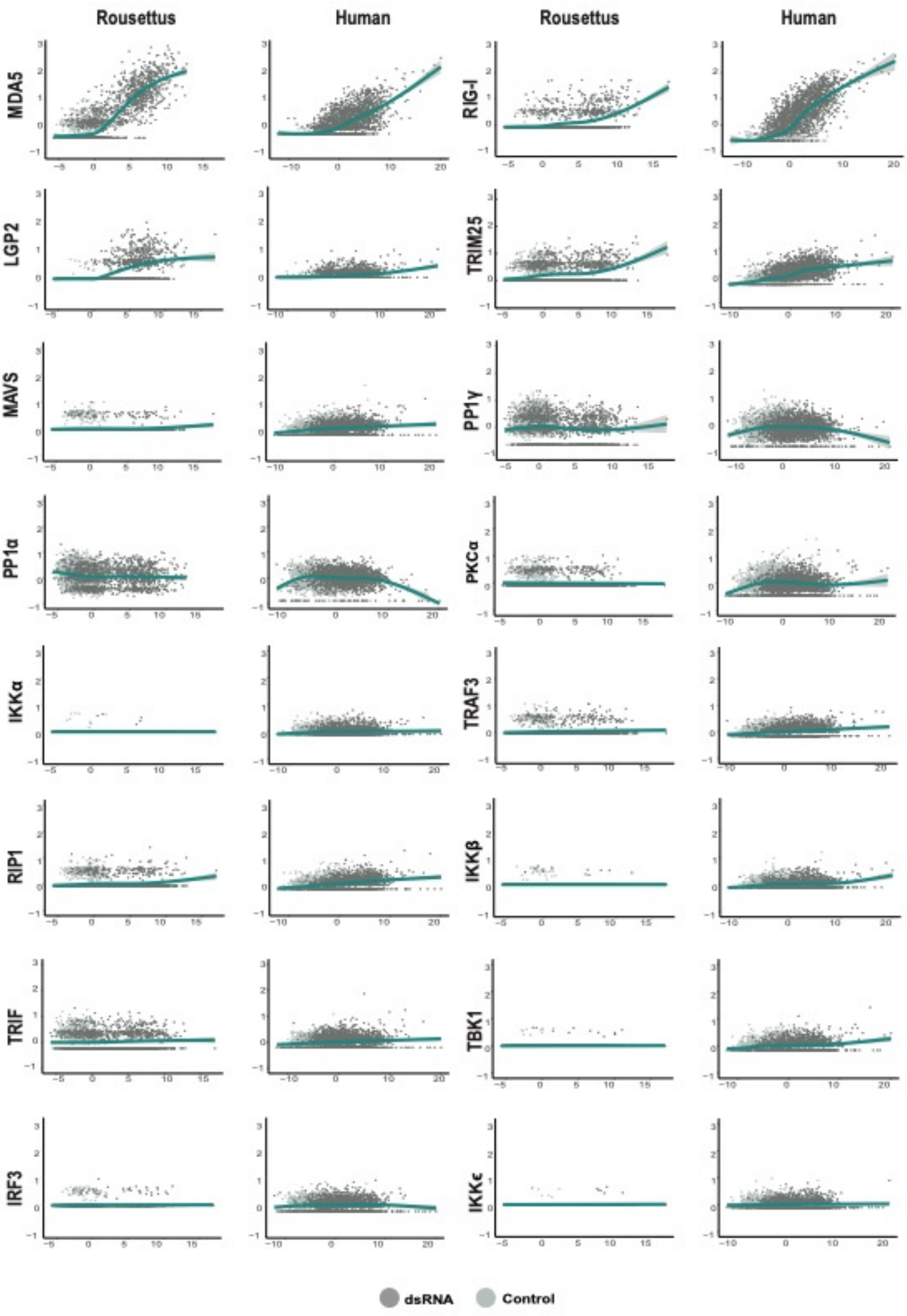


**Supporting Figure 7: Pseudotime analysis of genes in the RLR and TLR pathways, based on single-cell data of human and *Rousettus* cells stimulated with dsRNA and control.** Pseudotime is based on PCA of all gene expressed, as explained in Methods. Human and *Rousettus* cells were stimulated and profiled in parallel.

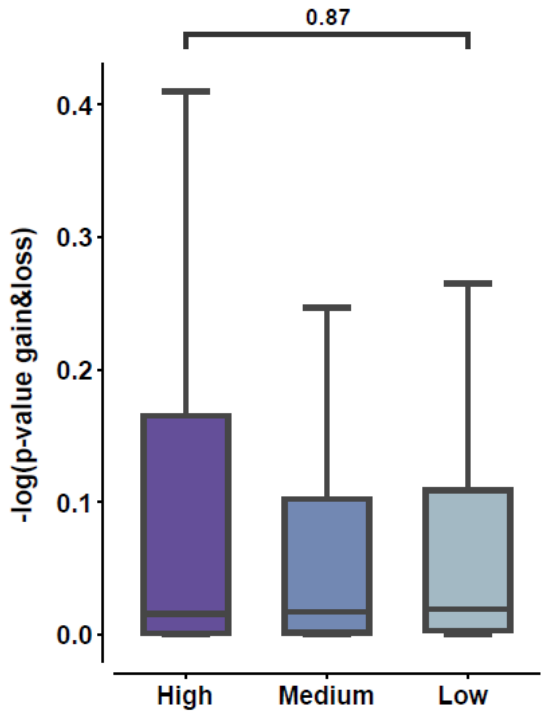


**Supporting Figure 8: Relationship between gene response divergence across species and gene's rate of gain and loss.** Genes are partitioned into three groups of low-, medium- and high- divergence in response to dsRNA between species, as explained in the main text. The distribution of P-values indicative of gene's evolutionary rate of gain and loss across the vertebrate clade, is shown (with negative logarithm) for each of these three groups. Distributions of high- and low-divergence groups are compared using a Mann-Whitney test.

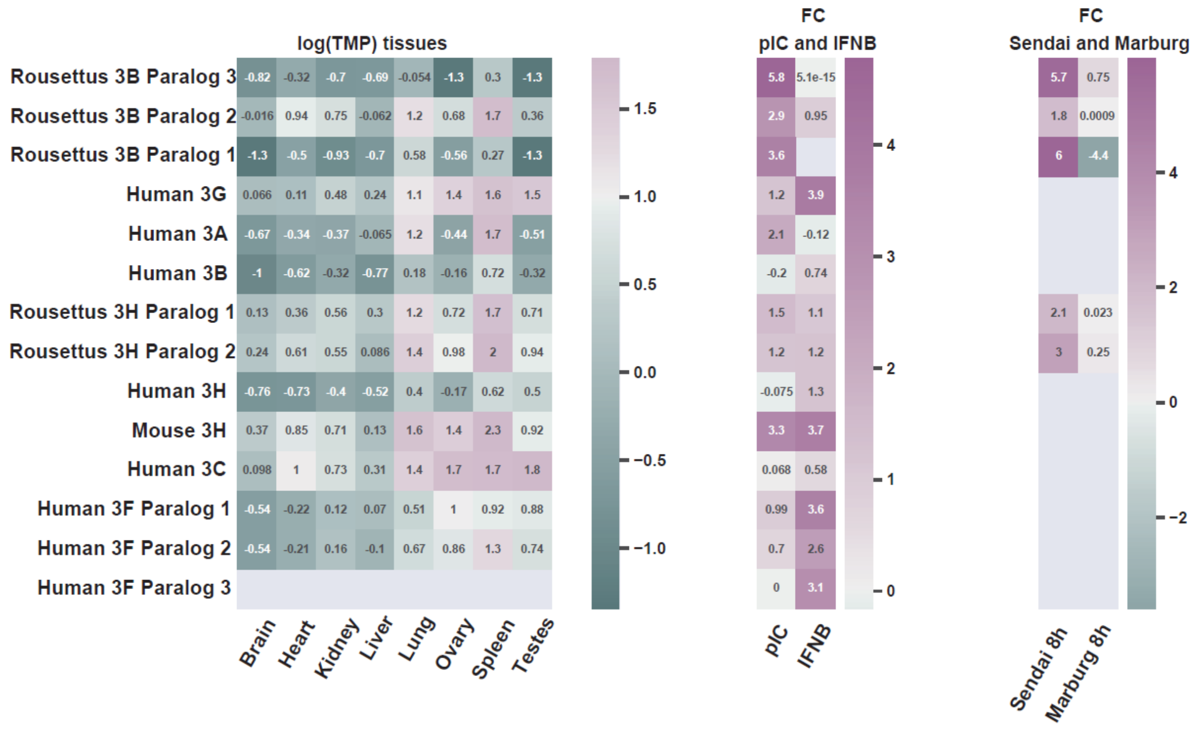


**Supporting Figure 9: Expression levels of APOBEC3 genes across eight tissues in human, mouse and *Rousettus*.** Genes were tested for their basal expression in eight equivalent tissues in human, mouse and *Rousettus*. Clade-specific duplicates are shown for *Rousettus* APOBEC3B and human APOBEC3H and 3F. The separated middle column shows each of these genes' fold-change in response to dsRNA and to IFN. The separated right column shows each of the Rousettus genes' fold-change in response to Sendai virus and Marburg virus infection. Genes that are not expressed appear in grey.
